# Tbc1d15 knockdown *in vivo* alleviates α-synuclein-induced neurotoxicity by promoting autophagy

**DOI:** 10.1101/2024.10.01.616109

**Authors:** F. H. Siddiqi, N. R. Raval, M. Palner, J. E. Nielsen, R. Pocock, D. C. Rubinsztein, P. Ejlerskov

## Abstract

Parkinson’s disease is a neurodegenerative disease, which is associated with accumulation of α-synuclein protein aggregates and Lewy Body formation. These neurotoxic inclusions are especially harmful for dopamine-producing neurons in the substantia nigra of the brain. The cellular degradation system autophagy can reduce neurotoxicity caused by accumulated α-synuclein, by targeting it for degradation. Previously, we demonstrated that human TBC1D15 inhibits autophagy *in vitro*, resulting in accumulation of neurotoxic protein aggregates. Conversely, lowering the TBC1D15 expression promotes autophagy and degradation of α-synuclein and huntingtin proteins in various cell models. Here we show that knockdown of murine Tbc1d15 *in vivo* activates autophagy, reduces α-synuclein-mediated neurotoxicity, and improves motor performance. Thus, targeting Tbc1d15 expression may be a therapeutic avenue for neurodegenerative diseases.

## Background

Parkinson’s disease (PD) is one the most common movement disorders that is also associated with cognitive decline^1^. A key finding in the brains of post-mortem PD patients is accumulation of α-synuclein, which forms neurotoxic oligomers and larger protein aggregates termed Lewy bodies^2^. Dopaminergic neurons in substantia nigra are especially vulnerable to α-synuclein accumulation, and animal models show gradual retraction of neurites that innervates the striatum, and loss of cells in the substantia nigra^3^. Excess α-synuclein is sufficient to cause PD, as autosomal dominant PD is caused by multiplication of the SNCA gene encoding α-synuclein, which increase its expression^4^.

Autophagy is cytoplasmic degradation system that breaks down cell organelles, long-lived proteins, lipids, invading pathogens, and neurotoxic protein aggregates by capturing them in double-membraned vesicles called autophagosomes. The autophagosomes ultimately deliver the substrates to lysosomes for degradation^5^. LC3-I is a cytosolic protein but upon autophagy activation it is conjugated to phosphatidylethanolamine and incorporated into nascent autophagosome membranes as LC3-II. LC3-II is associated with autophagosomes from the precursor stage until after fusion with lysosomes and LC3-II levels correlate with cellular autophagosome volumes^6^. The fusion step is orchestrated by the homotypic fusion and protein sorting (HOPS) complex, the syntaxin17-SNAP29-VAMP7/8 trans-SNARE complexes, and the Rab GTPase protein Rab7^7–10^.

TBC1D15 is a Rab GTP activating protein (GAP) that belongs to the TBC (Tre-2/Bub2/Cdc16) protein family. These proteins convert active GTP-bound Rab to inactive GDP-bound Rab^11^. We previously demonstrated that TBC1D15 inhibits autophagy by inactivating Rab7, resulting in the accumulation of autophagy substrates like the neurotoxic proteins α-synuclein and huntingtin^12,13^. We discovered that the microRNA miR1 targets and reduces TBC1D15 levels, which activates autophagy and lowers the neurotoxic burden from huntingtin and α-synuclein *in vitro*. Here, we show that shRNA targeting of murine Tbc1d15 *in vivo* promotes brain autophagy and protects dopaminergic neurons in substantia nigra from α-synuclein-induced neurotoxicity.

## Results

Measuring *in vivo* autophagy flux in the brain is challenging. Western blotting of brain extracts will only provide a steady state measurement, and it is not possible to determine whether an increase in LC3-II is caused by a blockage or an increase in autophagy. Hence, to validate our *in vitro* findings in a mammalian *in vivo* model, we used transgenic mice expressing mRFP-GFP-LC3, which enables *in vivo* monitoring of autophagy flux^14^. GFP and mRFP are fluorescent inside autophagosomes but in low-pH lysosomes the GFP signal is quenched. This enables quantification of autophagosomes, which are vesicular structures positive for GFP and mRFP, and autolysosomes which are only mRFP-positive vesicles. Thus, the number of mRFP-only vesicles reflects autophagy flux. mRFP-GFP-LC3 mice were injected in the substantia nigra with adeno-associated virus (AAV) containing shRNA targeting Tbc1d15 (left side) and scrambled control shRNA (right side) (Figure 1A). shRNA expression was regulated by the neuron-specific synapsin-1 promoter and was co-expressed with a cytosolic BFP reporter. BFP-positive neurons had a significant reduction in Tbc1d15 signal when compared to BFP-negative cells in the brain-half receiving shRNA against Tbc1d15 (Figure 1B-D, E). By contrast, no difference in Tbc1d15 signal was observed in BFP-positive neurons that received scrambled shRNA when compared to BFP-negative cells (Figure 1B-D, E). Furthermore, BFP-positive neurons (Tbc1d15 knockdown) had increased autophagy flux, as evidenced by an increase in both mRFP-positive vesicles and the mRFP/GFP ratio (Figure 2).

**Figure 1.**
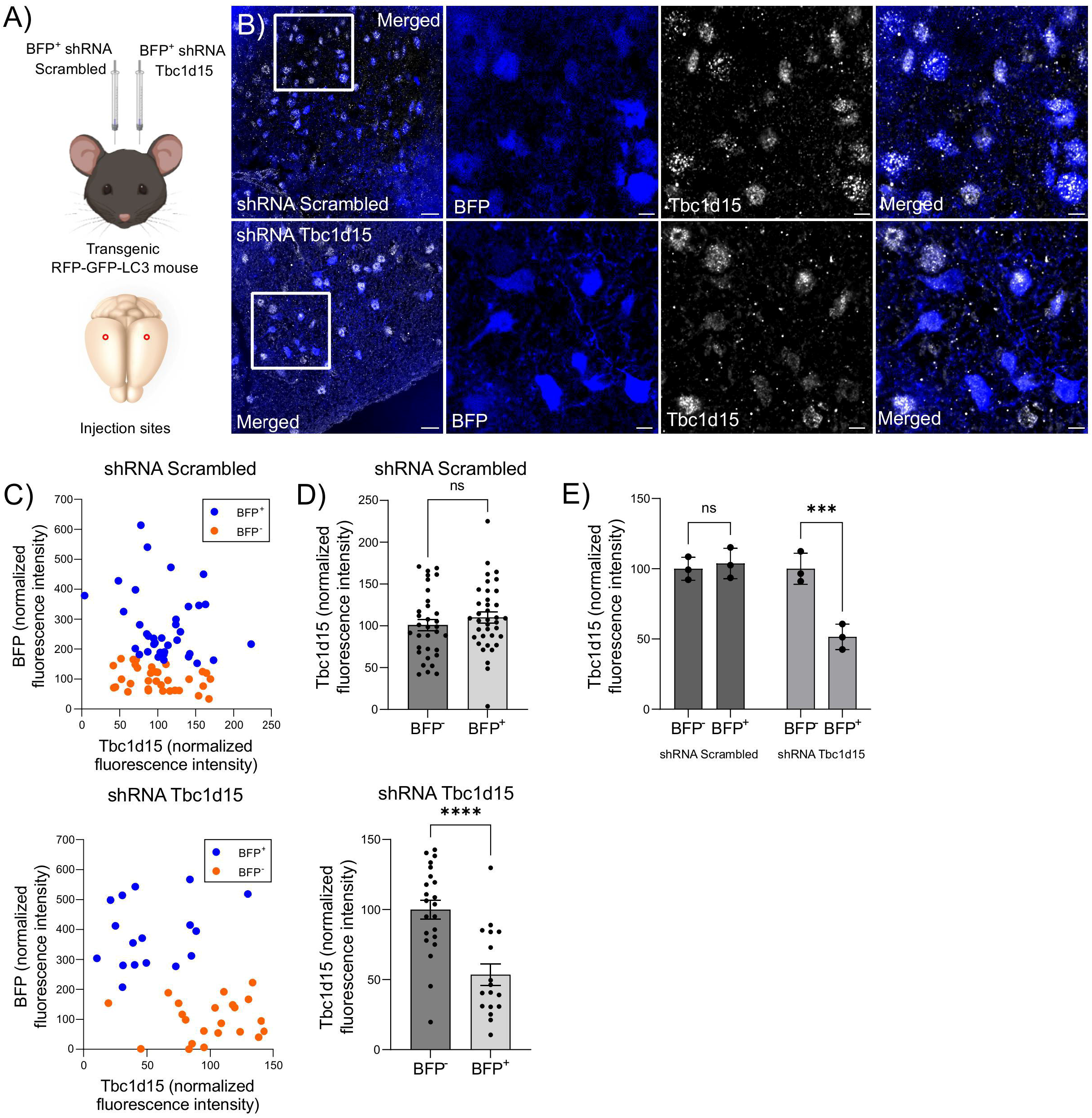
AAV shRNA reduces neuronal Tbc1d15 *in vivo*. 24-week-old transgenic mice expressing mRFP-GFP-LC3 were injected in the substantia nigra area with adeno-associated virus containing shRNA against Tbc1d15 (left side) or scrambled (right side). shRNA expression was regulated by the neuronal-specific promoter synapsin-1 and expresses a BFP reporter. 14 weeks after the injections the brains were PFA-perfused and processed for immunofluorescence imaging of the substantia nigra. A) Schematic of the AAV injection sites. B) Immunofluorescence images showing expression of Tbc1d15 in shRNA positive neurons indicated by the BFP-positive cells. Scale bar, 50μm; close-up, 10μm. C) Fluorescence intensity of Tbc1d15 and BFP in individual neurons, and D) mean Tbc1d15 fluorescence intensity in BFP-positive neurons obtained from the representative images in B). Student’s t-test, ****p = 0.0001, n = 39-70 individual cells from one image. E) Tbc1d15 mean fluorescence intensity in neurons from three individual mice. Quantified data is based on 5-10 images obtained from each side of mice, N = 3. ANOVA test, Šídák corrected, ***p = 0.001, ns = non-significant.

**Figure 2.**
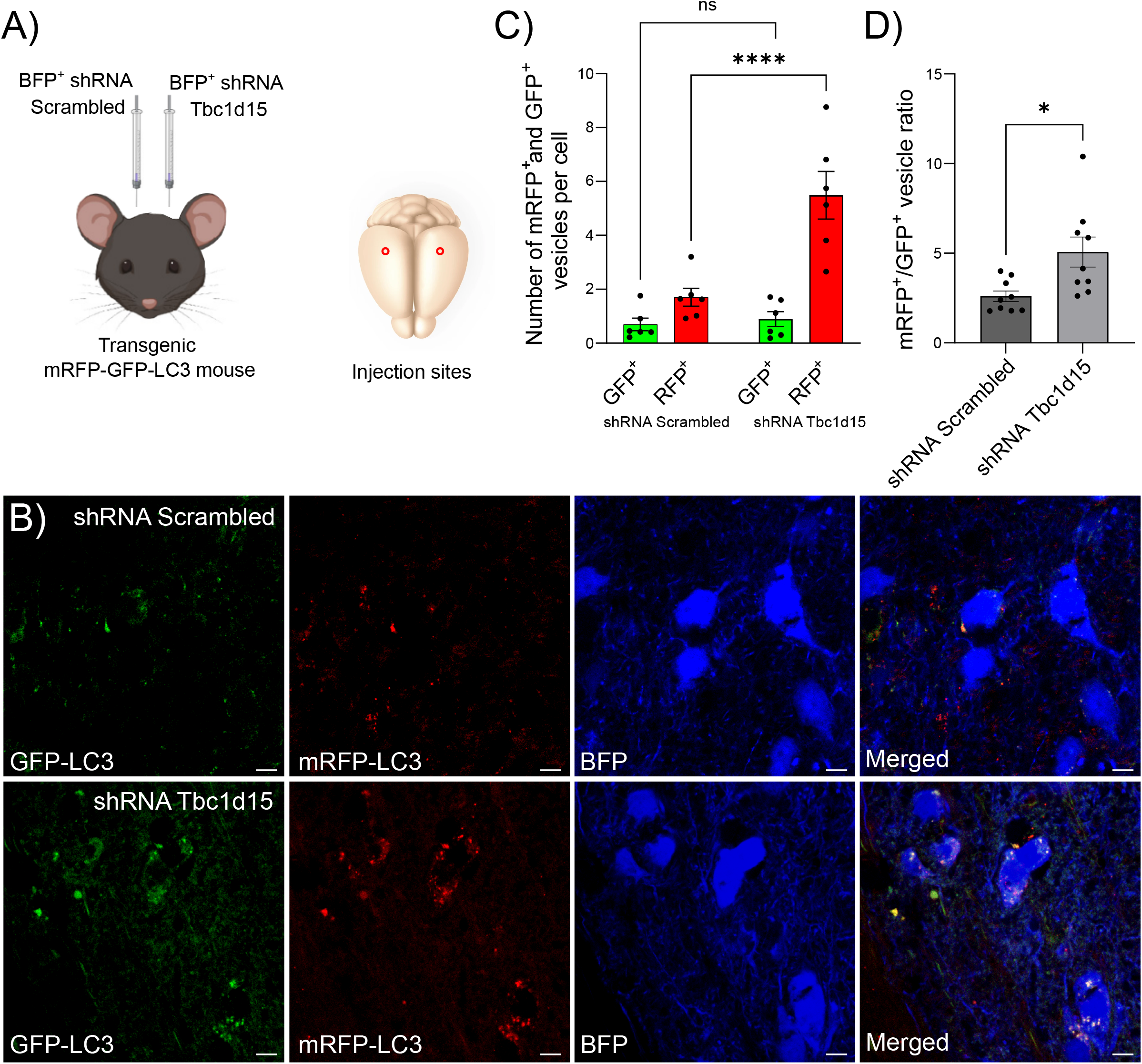
Tbc1d15 knockdown promotes neuronal autophagy *in vivo*. 24-week-old transgenic mice expressing mRFP-GFP-LC3 were injected in the substantia nigra area with adeno-associated virus containing shRNA against Tbc1d15 (left side) or scrambled (right side). shRNA expression was regulated by the neuronal-specific promoter synapsin-1 and expresses a BFP reporter. 14 weeks after the injections the brains were PFA-perfused and processed for confocal microscopy and quantification of LC3 positive vesicles in substantia nigra. A) Schematic of the AAV injection sites. B) Representative immunofluorescence images showing LC3 vesicles in shRNA-transduced neurons (BFP-positive). Scale bar, 10μm. C) Quantification of mRFP- and GFP-positive LC3 vesicles and D) the ratio between mRFP and GFP vesicles obtained from BFP-positive neurons in the substantia nigra. Quantified data is based on 7-15 images from each mouse, N = 9. C) ANOVA test, Šídák corrected, ****p < 0.0001; and D) Student’s t-test, two-tailed, *p < 0.05.

Next, we tested if Tbc1d15 knockdown could limit α-synucleinopathy *in vivo*. Eight-week-old Sprague Dawley rats were injected in the substantia nigra with AAV expressing human α-synuclein under the neuron-specific synapsin-1 promoter. In addition, shRNA against Tbc1d15 was injected in the left-side and scrambled shRNA in the right-side of substantia nigra (Figure 3A). Rat motor coordination was tested in a cylinder test both before and three weeks after the injections. The cylinder test evaluates the asymmetric behaviour of rodent front paws. When the rat explores the top of the cylinder, it uses its front paws for support on the cylinder wall, either with both front paws or with a single front paw (Figure 3B). The number of right- and left-front paw touches was counted over a 5-minute period. The front paws of the rat are controlled by the contralateral brain hemisphere, meaning that the right front paw is controlled by the left-brain hemisphere and the left front paw by the right-brain hemisphere. Three weeks of human α-synuclein expression in the substantia nigra has been shown to cause neurodegeneration of dopaminergic neurons, which affects motor coordination^15^. Prior to the α-synuclein overexpression, the rats used both front paws equally and most frequently with two-front paw stands (Figure 3C). After three weeks of α-synuclein overexpression, the rats used their right front paw more frequently, often as a single-front paw stand, suggesting more severe neurodegeneration in the right side of substantia nigra (Figure 3C-D). This suggests that Tbc1d15 knockdown in the left-side of the substantia nigra alleviated α-synuclein-induced motor defects.

**Figure 3.**
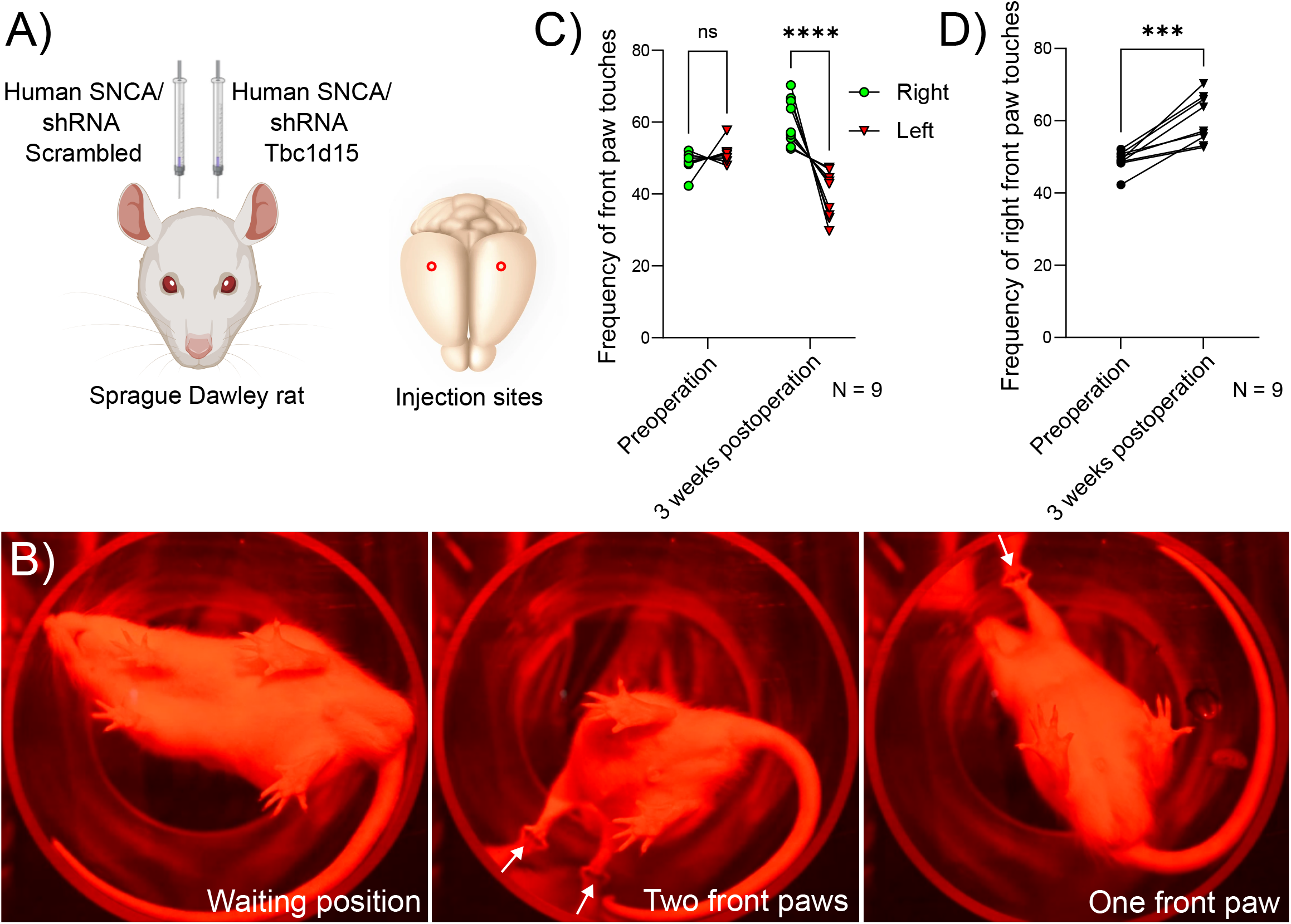
Tbc1d15 knockdown alleviates α-synuclein-induced behavioural defects. A) Eight-week-old wild-type Sprague Dawley rats were co-injected with adeno-associated virus expressing human α-synuclein (SNCA) (both sides of the brain) and shRNA against Tbc1d15 (left side) or scrambled (right side) in the substantia nigra. Transgenic expression was regulated by the neuron-specific promoter synapsin-1. B) Examples of behavioural phenotypes observed in the cylinder test. C) Frequency of front paw touches (left versus right) and D) right front paw touches pre-operation and three weeks after. C) ANOVA paired test, Šídák corrected, ns = non-significant, ****p < 0,0001, and D) paired Student’s t-test, two-tailed, ***p < 0.001.

Consistent with our behavioural data, we found that tyrosine hydroxylase expression was higher in the left-brain half that received shRNA against Tbc1d15, suggesting better preservation of dopaminergic neurons (Figure 4A-B). This was associated with a decrease in both Tbc1d15 and total α-synuclein levels (Figure 4A-B). The reduced presence of dopaminergic neurons was further validated by immunostaining of the brain, which revealed significantly reduced tyrosine hydroxylase immunoreactivity in both the neurite projections into striatum and in the substantia nigra (Figure 4C-E). Together, these data suggest that reducing Tbc1d15 expression is protective against α-synuclein-induced dopaminergic neurotoxicity.

**Figure 4.**
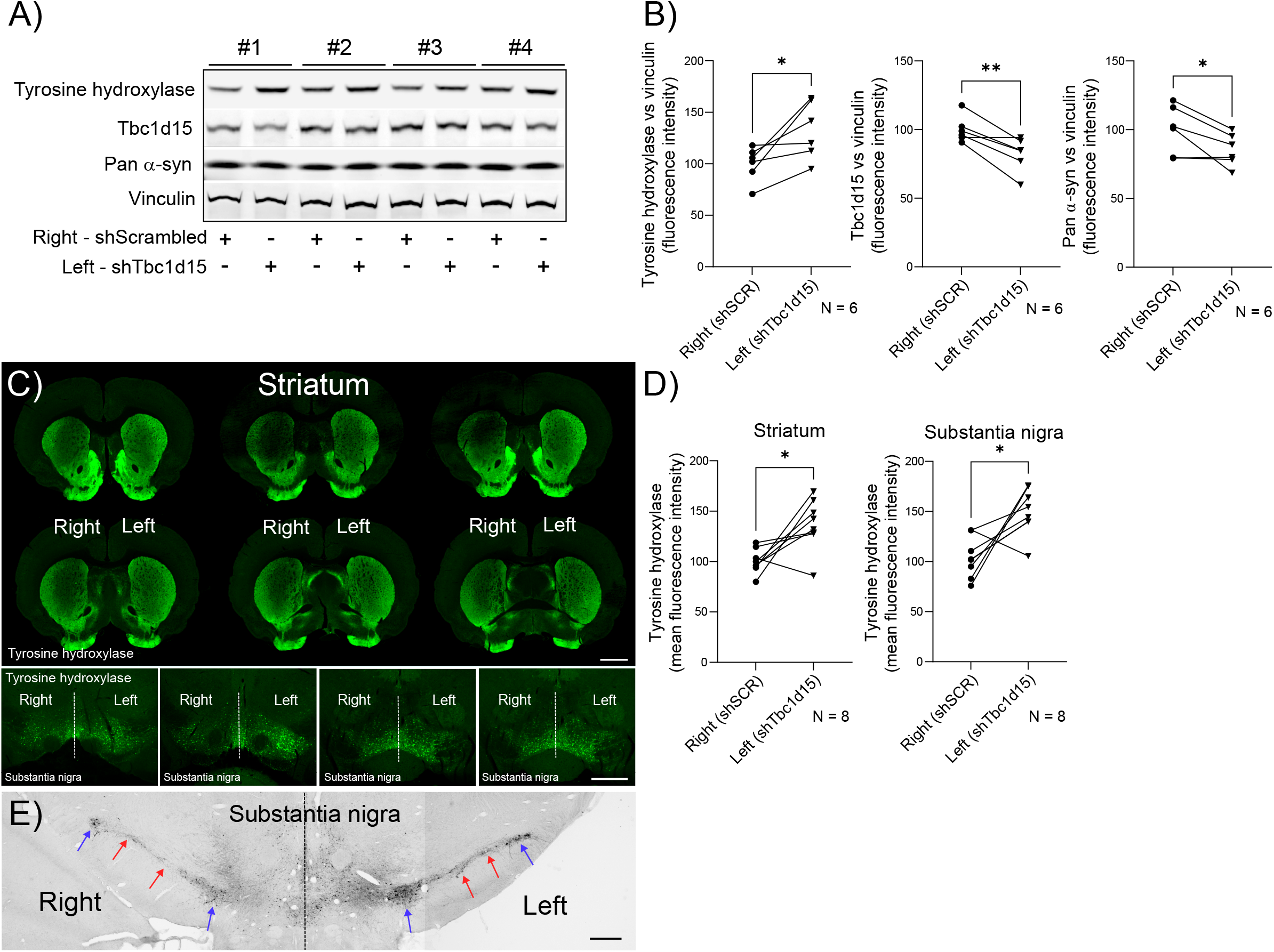
Tbc1d15 knockdown alleviates α-synuclein-induced neurotoxicity. A) Western blots of protein brain extracts (right vs. left brain hemisphere) from four individual rats and B) quantified protein levels (N = 6). Paired Student’s t-test, two-tailed, *p < 0.05, **p < 0.01. C) Upper panel; Immunofluorescence images of tyrosine hydroxylase staining in six consecutive brain sections covering the striatum of one mouse. Lower panel; tyrosine hydroxylase immunostaining of substantia nigra of four individual mice. Scale bar, 2mm. D) Quantified tyrosine hydroxylase immunofluorescence intensity comparing the two brain hemispheres in striatum and substantia nigra. Paired Student’s t-test, two-tailed, *p < 0.05, N = 8. E) Immuno-DAB (3, 3’-diaminobenzidine) staining against tyrosine hydroxylase showing fewer cell bodies (blue arrows) and neurite projections (red arrows) in the right substantia nigra region. The image is composed of three stitched images. Scale bar, 500μm.

## Discussion and conclusion

Here we show that reducing murine Tbc1d15 expression *in vivo* decreases α-synuclein-induced neurotoxicity. Previously we discovered that human TBC1D15 is negatively regulated by the microRNA, miR1, which increases upon interferon-β treatment^13^. Interferon-β, a type I interferon, is upregulated upon viral infection to elicit an anti-viral immune response^16^. Viral infections are detected by the cyclic GMP–AMP synthase (cGAS), which activates the stimulator of interferon genes (STING) pathway, that then promotes activity of the transcription factor IRF3^17^. IRF3 upregulates type I interferons, which bind to the interferon α/β receptor (IFNAR) on cell surfaces. This triggers the phosphorylation of STAT transcription factors, which homo- and heterodimerize and initiate a full anti-viral response^18^. Previously, we discovered that mice lacking the IFN-β gene exhibit pathogenic hallmarks of PD, such as neuromuscular and cognitive decline, as well as accumulation of neurotoxic α-synuclein^19^. This was due to a blockage in autophagy. Conversely, treating neurons with recombinant IFN-β activated autophagy and prevented α-synuclein-induced neurotoxicity. Hence, we have provided *in vitro* and *in vivo* evidence that IFN-β activates autophagy via a miR1-TBC1D15-Rab7 axis, and that reducing TBC1D15 levels provides a therapeutic strategy for neurodegenerative proteinopathies.

TBC1D15 has also been shown to be involved in mitophagy. Parkin and PTEN induced kinase 1 (PINK1) regulate mitophagy, which is the selective autophagy of mitochondria, and mutations in the parkin and PINK1 genes have been linked to PD^20^. Parkin also mediates amino acid flux between lysosomes and mitochondria by providing non-degradative contact sites between the two organelles^21^. This process is dependent on active Rab7, and the parkin mutant C431S destabilizes active Rab7. Intriguingly, lowering TBC1D15 in dopaminergic neurons generated from induced pluripotent stem cells (iPSC) from PD patients (parkin C431S), restored lysosomal-mitochondrial dynamics by facilitating Rab7-dependent mitochondrial and lysosomal tethering^21^.

During mitophagy, Atg9-positive membranes are recruited to mitochondria and contribute with membrane to the autophagosomal phagophore that engulf the mitochondria^22^. The recruitment of Atg9 positive vesicles is dependent on active Rab7a^23^. Compellingly, knocking out the two GAPs TBC1D15 and TBC1D17 enhances the co-localization between Atg9 and LC3-positive autophagosomes^23^, and coherently, TBC1D15 knockout cells accumulate LC3-positive structures upon mitochondrial damage^24^.

TBC1D15 contains a LC3-interacting region (LIR) that interacts with the Atg8 proteins GABARAP, GABARAPL1 and -L2, and can bind to Fis1 at mitochondria^25^. Thus, TBC1D15 has the potential to bridge mitochondria to the phagophore. During valinomycin-induced mitochondrial damage, which triggers parkin-mediated mitophagy, TBC1D15 is recruited to mitochondria and Rab7 associates with the phagophore^24^. In contrast to wild type TBC1D15, mutant TBC1D15 (D397A), which lacks TBC domain GAP activity, was unable to restore autophagy upon mitochondrial damage in TBC1D15 knockout cells. This suggests that TBC1D15 inactivates Rab7 to promote mitophagy of damaged mitochondria, which seems contradictory to the above mentioned results.

Recently, TBC1D15 was discovered to also play a role in lysosomal membrane repair^26^. Small perforations in the lysosome membrane are repaired by the ESCRT complex^27^, whereas more severe damage initiates lysophagy, the selective autophagy of lysosomes^28^. If the damage can be repaired, TBC1D15 is recruited to the damaged lysosome where it serves as a scaffold for assembly of the autophagic-mediated lysosomal reformation machinery. TBC1D15 depletes active Rab7 from the site, which may impede lysophagy, and instead drive autophagic-mediated repair of the damaged lysosome. A similar mechanism may occur in valinomycin-induced mitochondrial damage^24^, where the recruitment of TBC1D15, Rab7, and the phagophore could indicate an autophagosome-mediated membrane repair mechanism, instead of mitophagy. However, this requires further investigation.

The roles of TBC1D15 in autophagy and membrane repair mechanisms appear context dependent. Although removing TBC1D15’s inhibitory function on Rab7 is neuroprotective in the context of neurotoxic proteins that cause neurodegenerative diseases, as demonstrated here, pharmacological targeting of TBC1D15 should be thoroughly tested in relation to mitophagy and lysosomal repair mechanisms. Hence, a moderate inactivation of TBC1D15 could be a preferable strategy for potential pharmacological intervention.

## Materials and Methods

### Experimental animals

Mice were housed in individually ventilated cages with free access to standard animal food chow and water, in a climate-controlled room (45-65 % humidity and 20-24 ºC room temperature) with a 12 h light/dark cycle. Mouse studies were performed in accordance with the UK Animals (Scientific Procedures) Act with appropriate Home Office Project and Personal animal licenses and with University of Cambridge Ethics Committee approval. Mice were maintained and used in experiments following ARRIVE guidelines.

The Sprague Dawley rats were housed under standard laboratory conditions with 12-h light/dark cycles and *ad libitum* access to food and water. All experiments conformed to the European Commission’s Directive 2010/63/EU with approval from the Danish Council of Animal Ethics (Journal No. 2017-15-0201-01375) and the Department of Experimental Medicine, University of Copenhagen.

### Mouse model

To measure the effect of Tbc1d15 knockdown and *in vivo* upregulation of autophagy, we used a transgenic mRFP-GFP-LC3 autophagy reporter mouse line. The mRFP-GFP-LC3 mouse line was generated in the Rubinsztein lab^29^ and has been extensively used^30,31^. Mouse litters were produced by mating of transgenic mice and were genotyped at an age of 3 weeks by PCR using ear punches by Transnetyx (transnetyx.com). For mRFP-GFP-LC3 mice, mRFP probe, forward primer: AGCGCGTGATGAACTTCGA, and reverse primer: GCGCAGCTTCACCTTGTAGAT, were used by Transnetyx.

### Injection of adeno-associated virus (AAV) into the substantia nigra

For substantia nigra brain injections, 24-week-old mRFP-GFP-LC3 (males and females) were anesthetized via continuous administration of isoflurane (gas). The head was shaved, and the skin disinfected with Hibiscrub. Fully anesthetized mice were placed on a stereotactic apparatus and the front teeth were latched on the anterior clamp of the gas anaesthesia mask. After securing the mouse head on a stereotaxic frame, a 1.0 cm long midline linear incision was made on the skull from the frontal cranial bones to the back of the parietal cranial bones. The skin was pushed aside, the periosteum was removed using sterilised cotton buds so that the skull became visible. The bregma on the skull was identified, and the bregma and lambda were aligned on the same horizontal position. The target coordinate was marked for the substantia nigra region of the right and left hemispheres of the brain; anteroposterior (AP): -3.0mm; mediolateral (ML): ±1.3mm; dorsoventral (DV); −4.4mm relative to bregma), as described^32,33^. The holes on the right and left hemispheres were drilled into the skull using a dental micro drill. The needle (33 gauge, Hamilton 65461-01) was inserted into the brain through the craniotomies at the target coordinates. 2μl of scramble shRNA was injected on right hemisphere and 2μl of Tbc1d15 shRNA in the left hemisphere of the brain. The injections/infusions were performed at a constant rate of 200nl per minute using a syringe pump. The needle was left in place for 6 min to minimize upward flow of solution. The scalps were then sutured, and the mice were recovered and buprenorphine administered for post-operative analgesia. After 14 weeks, mice (38 weeks old) were sacrificed and brains harvested.

A similar approached were used for rats with the following modifications. Eight-week-old wild type Sprague Dawley rats were co-injected with AAV expressing human α-synuclein (SNCA) (both sides) and shRNA against Tbc1d15 (left side) or scrambled (right side) of the substantia nigra using a stereotaxic frame. The following coordinates were used relative to bregma: AP: -5.4 mm; ML: ± 1.7 mm; DV: -7.2 mm from below dura (−7.8 mm from skull surface). Holes were drilled in the scull with a diamond bore (diameter 0.5 mm). A 33 g needle (World Precision Instruments, Sarasota, FL, United States) was lowered (1 mm per 10 seconds) and 5 ul AAV solution injected at a speed of 275nl per minute. After 6 minutes the needle was retracted (1mm per 15 seconds).

### Adeno-associated viral vectors

All AAV vectors were designed, and viruses generated by VectorBuilder. All vectors were regulated by transgene expression under the neuron-specific synapsin-1 promoter and were produced in AAV serotype 9. SMART pool shRNA against mouse or rat Tbc1d15 (or scrambled) was designed in a miR30-based vector system. For the mouse-specific shRNA, a cytosolic BFP reporter was included in the vector backbone, to enable detection of neurons positive for shRNA when performing confocal microscopy imaging.

### Behaviour

Rat motor coordination was tested in a cylinder test before and three weeks after the AAV injections. Each rat was placed in a transparent cylinder (diameter: 10 cm; height: 40 cm) and behaviour recorded from below. When the rat explores the top of the cylinder it uses the front paws for support on the cylinder wall. The number of right and left front paw touches were counted over a 5-minute period to determine asymmetric motor behaviour. The front paws are controlled by the contralateral brain hemisphere, which means that the right front paw is controlled by the left-brain hemisphere and the left front paw by the right brain hemisphere.

### Brain perfusion and Immunofluorescence

Animals (mRFP-GFP-LC3 mice (N = 10) and rats (N = 8)) were injected intra peritoneum with pentobarbital sodium (euthatal) for deep anaesthesia. The animals were then cardiac perfused with ice-cold phosphate buffered saline (PBS^-^) followed by 4% paraformaldehyde (PFA, pH 7.4) with a flow rate of 1 ml per minute. Brains were dissected from the scull and placed in 4% PFA overnight at 4°C. The next day, brains were dehydrated in 15% sucrose for 5 hours and then transferred to 30% sucrose overnight at 4°C for cryoprotection. The brains were then attached a sectioning plate with OCT (optimal cutting temperature) cryo embedding matrix, snap-frozen in crusted dry-ice, and thin sections were prepared with a cryostat.

Quantification of mRFP-GFP-LC3 vesicles in mRFP-GFP-LC3 mouse brains was carried out as previously described^30,31^. Briefly, 6μm thick coronal sections were cut from PFA perfused and cryo-protected mRFP-GFP-LC3 mouse brains. Sections were mounted on poly-lysine coated slides and air-dried for 2 hours at room temperature. To reduce autofluorescence, sections were stained in 0.05% Sudan Black in 70% methanol for 5-10 minutes^34^ and washed three time in PBS^-^. Sections were air-dried for 10 minutes and mounted using ProLong™ Gold Antifade (Life Technologies).

For each mouse, 5-12 images were acquired from the substantia nigra. The number of mRFP-GFP-LC3 dots (autophagosomes; mRFP and GFP-positive vesicles) and mRFP-LC3 dots (autolysosomes, only mRFP-positive vesicles) were counted in BFP-positive neurons using the Image J plugin ComDot. The experimenter was blinded to the identity of the samples.

For immunostaining of Tbc1d15, sections were incubated on Sudan Black as above and washed three times in PBS^-^, and then blocked in 5% normal goat serum, 2% bovine serum albumin, and 0.25% TX-100 in PBS^-^ (blocking buffer) for 45 minutes. The slides were incubated with primary antibody against Tbc1d15 (Abcam, cat. no. ab121396) in blocking buffer overnight. The following day the slides were washed three time in PBS^-^ and incubated with anti-rabbit secondary antibody Alexa fluor 633 and mounted with ProLong™ Gold Antifade. For qualification, masks were placed around BFP-positive neurons and BFP-negative cells, and the Tbc1d15 fluorescence intensity was quantified in the individual cells using region of interests (ROI) in Image J. Images of the mRFP-GFP-LC3 mice were acquired with a confocal LSM 810 (Carl Zeiss) confocal microscope or a Nikon Ti2 Crest X-Light V3 DeepSIM super-resolution spinning disc confocal using 405 nm, 488 nm, 568 nm, and 633 nm lasers.

For tyrosine hydroxylase staining, the whole rat brain was cut coronally at 35mm thick sections and transferred into anti-freeze medium (Na_2_HPO_4_(1.744g), NaH_2_PO_4_ (0.502g), ethylenglycerol (96ml), glycerol (96ml), dH_2_O (128ml). Free floating sections were immunostained with antibodies against tyrosine hydroxylase (Merck; cat. AB152) following the same procedure as above but using anti-rabbit secondary antibody Alexa fluor 488. Images were acquired in a LI-COR M Odyssey fluorescence scanner for the striatum and a EVOS M7000 microscope for the substantia nigra. Tyrosine hydroxylase fluorescence intensities were measured in masks placed around the striatum and substantia nigra in Image J. For immune-DAB (3, 3’-diaminobenzidine) staining of tyrosine hydroxylase the VECTASTAIN® ABC systems (Vector Laboratories, cat. no. PK-8200) was used according to the manufacturers protocol.

### Western blotting

Rat brains (n=6) were dissected and divided at the midline into right and left brain hemispheres, then snap-frozen in liquid nitrogen and stored at -80°C. Proteins were extracted in a dounce homogenizer containing TX-100 lysis buffer (1% TX100, 5M NaCl, 1M Tris-HCl, 10mM MgCl_2_, 1mM EDTA) using 2 times 20 strokes. Cell homogenates were centrifuges at 17,000g for 10 minutes at 4°C. The supernatants were collected, and the pellets centrifuged an additional round at 17,000 g for 10 minutes at 4°C, and the second supernatant collected. The protein concentration of the TX100-soluble fraction was measured with Dc Protein Assay (Biorad cat. no. 500-0116), added Laemmli buffer, and boiled at 100°C for 5 minutes. The samples were loaded (20-40 μg per well) on pre-casted 4-12% gels (cat. no. NW04125BOX) and for LC3 detection 16% gels were used (XP00165BOX). Proteins were separated by electrophoresis and transferred to pre-activated PVDF membranes (Millipore). The membranes were blocked in 5% milk or 5% BSA, and for α-synuclein the membranes were fixed in 0.4% PFA for 20 minutes prior to 40 minutes in 5% BSA. Membranes were incubated shaking overnight with primary antibodies (AbCam: Tbc1d15, cat. no. ab121396 and α-tubulin, cat. no. ab7291; SIGMA: vinculin cat. no. V9131; Cell Signaling Technology: α-synuclein cat. no. #2642; Novus Biologicals: LC3, cat. no. NB600-1384). The next day membranes were washed 3 times in PBS-T (PBS^-^ with 0.5% Tween-20) and incubated with secondary fluorescent antibodies (LiCor: IRDye® 680LT (H + L) and IRDye® 800CW Goat anti-Mouse and -rabbit IgG (H + L) shaking for 1 hour at room temperature. Finally, the membranes were washed 3 times in PBS-T and imaged in a LI-COR M Odyssey fluorescence scanner.

### Statistical analysis

The following statistical analysis were used: Student’s t-test pair and non-paired, two-tailed, and ANOVA with Šídák correction for multiple comparisons.

## Acknowledgements

We are grateful for funding from National Foundation for Medical Research and Innovation (Australia), Parkinsonforeningen (Denmark), Sundhedsdonationer (Denmark), the Nordic Autophagy Society, and the UK Dementia Research Institute, which is principally funded by the Medical Research Council.

